# Nearly half of all bacterial gene families are biased toward specific chromosomal positions

**DOI:** 10.1101/2023.10.18.562889

**Authors:** Xiao-Pan Hu, Martin J. Lercher

**Author notes:** Corresponding author (MJL).

## Abstract

Many bacterial genes encoding components of the translation and transcription machinery occupy genomic positions close to the origin of genome replication. These positional biases are thought to result from selection for high expression levels, as genes close to the origin are the first to be duplicated after the initiation of DNA replication. However, recent work indicates that positional biases of RNA genes involved in translation have evolved to support optimal growth rate dependence of expression levels rather than high expression per se. We hypothesized that, more generally, natural selection may have favored the location of different genes at specific chromosomal positions to optimize the growth rate dependence of their relative gene copy numbers. Here we show that 49% of bacterial gene families are preferentially localized to specific chromosomal regions, with most families biased toward either the origin or the terminus of replication. From our hypothesis, we derive six specific predictions regarding the genes’ chromosomal positioning and functional categories, as well as the minimum cellular doubling times. All six predictions are robustly supported by comparative genomic analyses of 773 bacterial species and by proteomics data from *Escherichia coli* and *Bacillus subtilis*. Our findings reveal a complex relationship between bacterial growth, resource allocation and genome organization, and they provide new insights into the physiological significance and potential functions of a large number of gene families.

## Introduction

The positioning of genes along the chromosome is a central aspect of genomic organization in bacteria^1–4^, with some gene families known to be biased toward particular chromosomal regions across bacterial species. For example, highly expressed genes tend to be located near the origin of replication (*oriC*)^4,5^; the exponential phase nucleoid-associated proteins and their targets are also located near *oriC* in *E. coli*^6,7^; and Spo0F and KinA, which together regulate sporulation in *B. subtilis*^8^, are located on opposite sides of the circular chromosome. However, we still lack a systematic understanding of the extent and the consequences of biases in gene positions across bacterial chromosomes^1^.

Gene positions influence transient, periodic variations in gene copy number due to replication-associated gene dosage^1,9^. Bacterial DNA replication begins at *oriC* and proceeds bidirectionally toward the terminus of replication (*ter*). During replication, genes already passed by the replication fork are present in more copies than genes still waiting to be replicated. This effect leads to a correlation between the expression patterns of genes along the two arms along the cell cycle at the single cell level^10^. At the population level, this process results in higher gene dosage for genes near *oriC* compared to those near *ter*. This effect is thought to be responsible for the preferential positioning of highly expressed genes near *oriC*^4,11^. Less appreciated is the dependence of replication-associated gene dosage effects on the growth rate^3^. At the lowest growth rates, replication occupies only a small fraction of the cell cycle, and the transient copy number changes have little effect. With increasing growth rate, however, the relative gene dosage (gene copy per genome equivalent of DNA) and hence the relative expression level of genes near *oriC* increases, while those of genes near *ter* decrease^12,13^ (**Fig. 1**). With the exception of functional RNA genes^14^ and sporulation genes in *B. subtilis*^8^, this growth rate dependence has not been studied systematically.

**Fig. 1.**
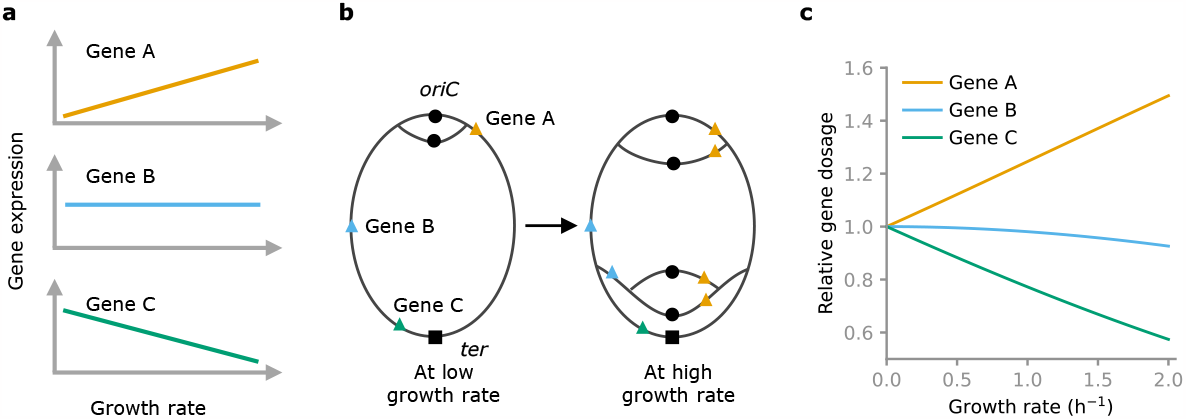
Scaling of gene expression with growth rate through replication-associated gene dosage effects depends on genomic gene positions. **(a)** The growth rate dependence of the expression of gene A (top), gene B (middle), and gene C (bottom). **(b)** At low growth rates (left), gene A, gene B, and gene C have the same chromosomal copy numbers throughout most of the cell cycle. At high growth rates (right), the copy number of gene A (at *oriC*) is higher than that of gene B (at *ter*) throughout most of the cell cycle. **(c)** As a result, the relative dosage of gene A increases with growth rate, while that of gene C decreases. Relative gene dosages are calculated in *E. coli* at chromosomal positions 0.15 (gene A), 0.5 (gene B), and 0.85 (gene C), see **Methods**. If the curves in (a) represent optimal expression patterns, then natural selection should favor the localization of gene A near *oriC* and of gene B near *ter* compared to other chromosomal localizations.

Replication-associated gene dosage effects provide a reliable, general mechanism that regulates gene expression as a monotonic function of growth rate. Growth rate-dependent gene expression, even independent of environmental details, has been observed for large sets of genes^15,16^. In particular, the expression of ribosomal genes in *E. coli* increases with growth rate^15^, while that of flagellar genes decreases^17,18^. The positions of these genes are consistent with growth rate-dependent regulation through replication-associated gene dosage effects (**Fig. S1**), with the genes encoding ribosomal proteins mostly located near *oriC*, leading to an increase of relative gene dosage with growth rate^3^, and flagellar genes located near *ter*, leading to a decrease of relative gene dosage with growth rate. That rRNA genes are closer to *oriC* than tRNA genes is also consistent with their relative expression, as the rRNA/tRNA expression ratio increases with growth rate across species^14^. In *E. coli*, average gene positions of rRNA and of tRNA genes are such that at the highest growth rate, their relative expression matches exactly their relative chromosomal copy numbers^14^. Finally, some cellular decisions – such as the transition to sporulation in *Bacillus subtilis*^8^ – are regulated through the growth rate-dependent balance of copy numbers between genes located at different chromosomal positions. Taken together, these observations suggest that at least some genes are located at specific chromosomal positions to support their optimal, growth rate-dependent expression rather than to favor high expression *per se*.

We hypothesized that more generally, natural selection may have favored the localization of genes to specific chromosomal positions in order to coordinate their expression across growth rates. This hypothesis leads to six specific predictions, which are detailed and tested below. Briefly, they concern (1) the extent of chromosomal position biases across gene families; (2) the distribution of positions along the chromosome preferred by different gene families; (3) how protein abundances of gene families biased to different chromosomal positions change with growth rate; (4) the extent of positional biases in a given genome as a function of its maximum growth rate; (5) the persistence of positional biases over vast evolutionary distances; and (6) the systematic variation of positional biases across functional gene categories.

## Results and discussion

### 49% of gene families are preferentially located at specific chromosomal positions

If natural selection has generally exploited replication-associated gene dosage effects to establish growth rate-dependent gene expression patterns, we expect to find many gene families that are biased toward specific chromosomal positions. We define a gene’s normalized chromosomal position (position for short) as its shortest distance to *oriC* along the circular chromosome, divided by half the chromosome length. Gene positions thus range from 0 (at *oriC*) to 1 (at *ter*) along both sides of the chromosome. We defined gene families such that each family corresponds to a single cluster of orthologous genes in the COG database^19^. With few exceptions, the COG dataset contains a single representative genome for each genus and thus largely eliminates potential biases caused by the unbalanced distribution of sequenced genomes across lineages (**Table S1**). To quantify the extent to which a given gene family is biased toward specific chromosomal locations, we first estimated the distribution peak (mode) of the corresponding gene positions (**Methods**). We then compared all windows of width 0.3 (corresponding to 30% of the complete genome) that contained this peak. The “position bias score” *b* was defined as the maximum fraction of genes from the family included in one of these windows (**Table S2;** see **Methods** and **Fig. S2**). Thus, if the genes of a given family are randomly positioned on bacterial chromosomes (not position-biased), its score *b* will be around 0.3; if all genes from a family are positioned close to the mode, the family is maximally biased, with a score *b* of 1. The distribution of position bias scores *b* across gene families is characterized by a sharp rise from just above 0.3 to 0.4, followed by a long tail toward 1.0 (**Fig. 2a**).

**Fig. 2.**
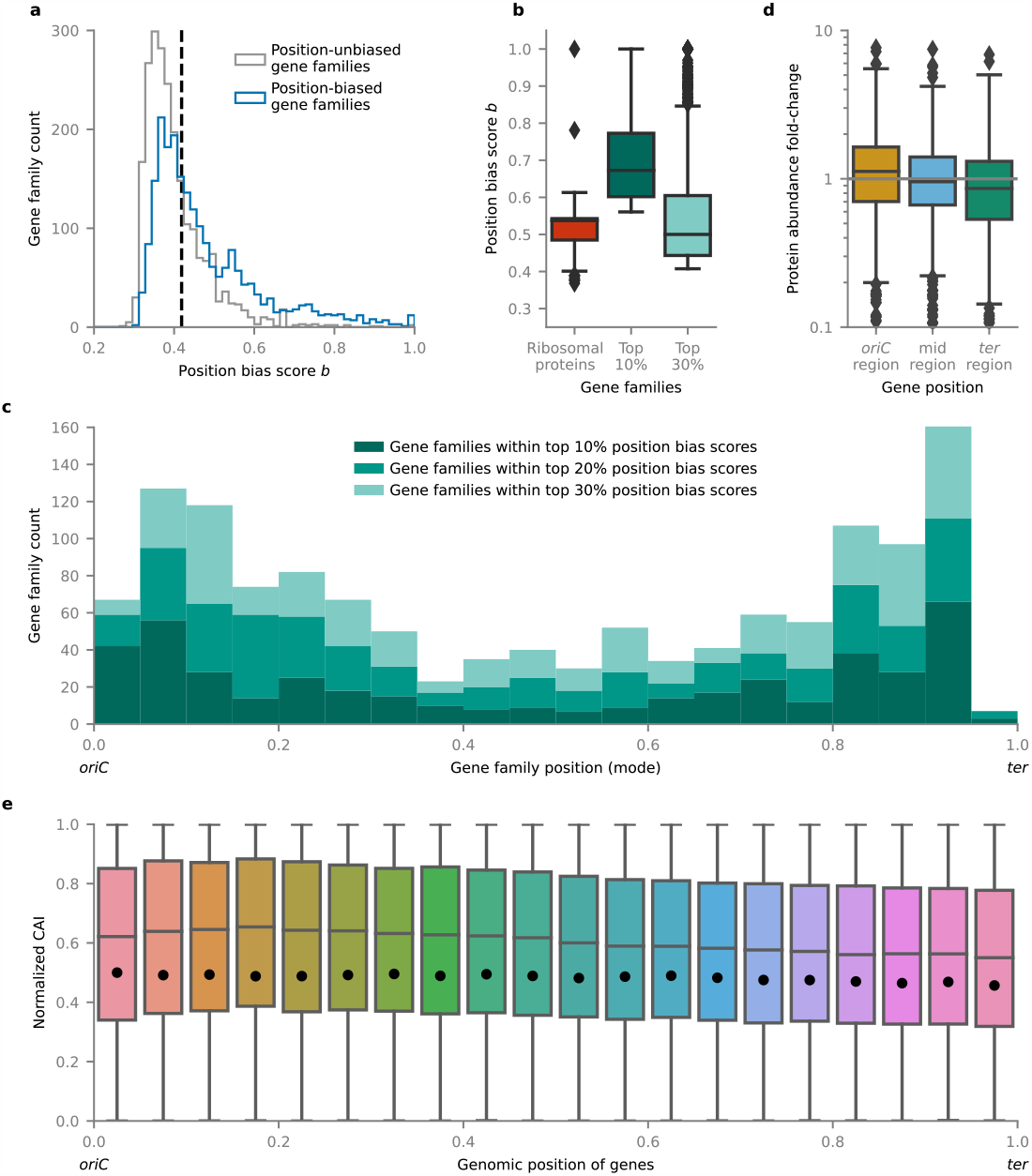
Distribution and expression of strongly position-biased genes. (**a**) Histograms showing the distributions of position bias scores of position-unbiased gene families and position-biased gene families. The vertical dashed line marks the threshold for strongly position-biased genes (*b*=0.4074), with the number of position-biased gene families above this threshold representing 30% of all gene families. (**b**) Position bias scores of genes encoding ribosomal proteins and other strongly position-biased genes. **(c)** Distribution of the mode (gene position peak) along the chromosome for strongly position-biased gene families. Colors mark position bias levels. **(d)** Changes in protein abundance in *E. coli* growing on rich medium (growth rate μ=1.93 h^-1^) vs. on a minimal medium (μ=0.74 h^-1^), quantified by ribosome profiling. Fold-changes of protein abundance are summarized for the products of all genes, divided into three equally sized chromosomal regions: 0-0.33 (*oriC*), 0.33-0.66 (mid), 0.66-1 (*ter*). **(e)** Normalized codon adaptation index (CAI) – a proxy for gene expression level – of strongly position-biased gene families (top 30% position bias scores) at different chromosomal positions. The dots mark the median of the normalized CAI of all other genes at the same chromosomal positions. In panels b, d and e, boxes extend from the first quartile to third quartile of the data; whiskers extend from the box to the last point included in 1.5x the interquartile range; lines inside the boxes mark the median; diamonds mark outliers.

We considered a gene family to be position-biased if there was statistically significant evidence for a non-random distribution of its gene positions (FDR-adjusted *p*-value < 0.05 from Kolmogorov-Smirnov test against uniform distribution, adjusted for multiple comparisons according to Benjamini-Hochberg^20^; **Methods**). Using this criterion, we identified 49.1% of the bacterial gene families in the COG database as position-biased (2177 out of 4430 gene families) (**Fig. 2a**). This number is orders of magnitude higher than previously reported estimates^4,6,21^.

Positional biases in bacterial genomes have so far been discussed mainly for ribosomal genes, which tend to be in close proximity to *oriC*^4,5,22^. Hundreds of bacterial gene families show much stronger positional biases (higher bias scores *b*) than most ribosomal genes: the top 10% most biased families (*n* = 443) contain only 6 out of 56 ribosomal genes (11%), two of which are found in only a minority of genomes (L14E in 7 genomes and L7Ae in 127 genomes). The bulk of bias scores for ribosomal genes is close to the median bias score of the top 30% most biased genes (**Fig. 2b**).

Note that if natural selection favors certain chromosomal positions for some gene families, the corresponding positions will be less accessible for genes from other families; these other families will therefore tend to be located in the remaining chromosomal positions, resulting in “passive” positional biases. Such passively biased gene families will generally have low position bias scores. To largely eliminate them from our analysis, we focus below on gene families with relatively high position bias scores, defined here as those in the top 30% (bias score *b* > 0.4074, *n* = 1329).

### Both *oriC* and *ter* regions are hotspots for strongly position-biased genes

Genes located near *oriC* and near *ter* experience substantial changes in relative gene copy number across growth rates, at least in fast-growing species (**Fig. 1**). This effect may have been exploited by natural selection in the case of genes for which growth rate-dependent changes of gene expression are selectively advantageous. Accordingly, we predict that the *oriC* and *ter* regions might be hotspots for strongly position-biased genes. As predicted, we found that the typical position (mode) of strongly position-biased gene families has two distribution peaks, located near *oriC* and near *ter* (**Fig. 2c**).

If these localizations are indeed caused by selection for the corresponding expression patterns, we expect the abundance of proteins encoded by genes near *oriC* to increase with growth rate, while those of proteins encoded by genes near *ter* should decrease with growth rate. This is indeed the case (**Fig. 2d**): the fold-change from slow growth (*μ*=0.74 h^-1^ on minimal medium) to fast growth (*μ*=1.93 h^-1^ on rich medium)^23^ of *E. coli* proteins is predominantly positive for those encoded near *oriC* (one-sided Wilcoxon signed-rank test: *p* = 8.5×10^−9^, *n* = 652); for those encoded near *ter*, the fold-change is predominantly negative (*p* = 2.4×10^−5^, *n* = 566). We confirmed these trends with another dataset in *E. coli*^24^ and in *B. subtilis*^25^ (**Fig. S3**). Thus, all the proteomics data examined is consistent with the hypothesis that natural selection for growth rate-dependent gene expression regulation drives positional biases on bacterial chromosomes.

Previous studies have postulated that the localization of ribosomal genes near *oriC* is related to natural selection for high levels of gene expression especially during rapid growth^4,5,22^ rather than growth rate-dependent regulation *per se*. To examine if strongly position-biased genes tend to be highly expressed in general, we used the codon adaptation index (CAI)^26^ as a proxy for gene expression. To facilitate the aggregation of results across genomes, we converted species-specific CAI values to ranks and normalized these to range from 0 (lowest CAI) to 1 (highest CAI; **Methods**). We found that strongly position-biased genes tend to be more highly expressed than other genes at the same genomic position not only near *oriC* – where replication increases the relative gene copy numbers – but throughout the bacterial chromosome (**Fig. 2e**; one-sided Wilcoxon rank-sum tests: *p* < 10^−15^ for each of the 20 bins). Since selection for high relative copy numbers can only act close to *oriC*, this finding supports our hypothesis that the selective force behind the establishment of position biases is related to growth rate-dependent gene regulation rather than to the gene expression levels themselves.

### Species whose DNA replication time exceeds their minimum cellular doubling time show the strongest positional biases

The time required for one round of DNA replication equals the speed of the DNA polymerase times the genome size. If this time is always much less than the cellular doubling time, replication-associated gene dosage effects are likely unimportant. In contrast, if the DNA replication time becomes comparable to the minimum cellular doubling time, genes near *oriC* will experience twice the dosage compared to genes near *ter* throughout most of the cell cycle. When DNA replication time exceeds the cellular doubling time, multiple simultaneous replication forks are required to synchronize DNA replication with cell growth, and dosage imbalances can double or quadruple. *E. coli* cells, for example, have been observed to contain up to eight origins of replication simultaneously^27^. Thus, if gene position biases indeed evolved to exploit replication-associated gene dosage effects for growth rate-dependent regulation of gene expression, we would expect to see the strongest gene position biases in species where the DNA replication time far exceeds the minimum cellular doubling time, and the weakest position biases in species where DNA replication times are small compared to their minimum doubling time.

To test these predictions, we calculated the ratio *R* between the estimated DNA replication time and the minimum cell doubling time^4^ for each species in our dataset. We were able to find estimates for minimum doubling time (maximum growth rate) for 416 of the 773 species in our dataset (**Methods**). These 416 species were divided into three groups: large-*R* species (*n*=139), with a high maximum number of replication forks; moderate-*R* species (*n*=139); and small-*R* species (*n*=138), with a low maximum number of replication forks (**Table S3**). We re-calculated the gene position bias scores and re-identified position-biased genes within each of the three groups, using the same procedure as applied previously to the full dataset. As predicted, the number of genes with statistically significant position biases differs systematically among the three groups (**Fig. 3a**). Gene families with high bias scores are most frequent in the large-*R* group and least frequent in the small-*R* group (**Fig. 3a** and **Table S2**): 1085 gene families in the large-*R* group have bias scores above 0.5, while this number is only 106 in the small-*R* group. That large-*R* species harbor more genes with strong position biases is not restricted to certain functional groups, but is found in all individual COG categories (**Fig. S4**). This latter finding contradicts a previous report that maximum growth rate affects the gene positions only of genes involved in translation and transcription^4^.

**Fig. 3.**
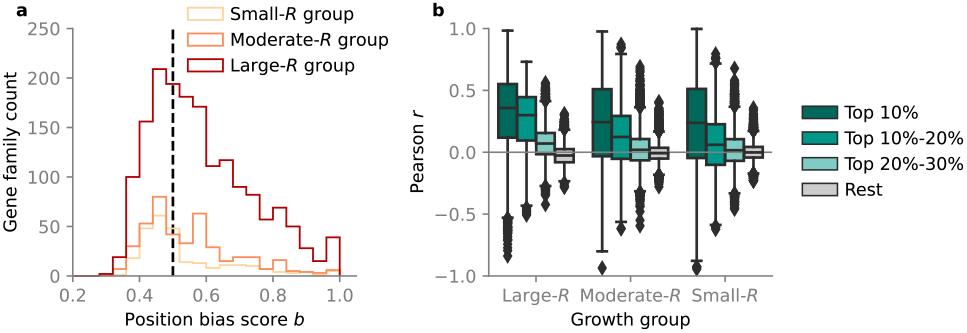
Species with long DNA replication times relative to the minimum cellular doubling time show the strongest positional biases. (**a**) Histograms showing the distributions of bias scores of position-biased genes in large-*R*, moderate-*R*, and small-*R* growth groups. The vertical dashed line marks position bias score *b*=0.5. (**b**) Distribution of Pearson’s correlation coefficient of gene positions for pairs of species from two different phyla, grouped by position bias scores of the corresponding gene families.

It is conceivable that positional biases arise from shared ancestry rather than from natural selection. In this case, one would expect little correlation between the positions of genes from the same gene family across large evolutionary distances. To test our hypothesis against this possibility, we compared the positions of genes across bacterial phyla, most of which diverged about 2 billion years ago^28^. Within each of the three *R* groups, we calculated Pearson’s correlation coefficient *r* of gene positions for families with different levels of position bias (top 10%; top 10%-20%; top 20%-30%; and all others). Note that for this analysis, the bias levels of gene families are those calculated from the entire dataset, not from the individual growth groups. For gene families without strong positional biases, the correlations of positions across phyla are distributed around zero (gray boxes in **Fig. 3b**). This finding supports the null hypothesis that in the absence of positional biases, genomic positions are effectively randomized over long evolutionary time scales. In contrast, all groups of strongly position-biased genes (top 10%, 10%-20%, 20%-30% of all genes, ranked by bias score *b*) show statistically significant positive correlations of gene positions across phyla in each of the three *R* groups (*p*<10^−15^ from one-sided Wilcoxon signed rank test for all cases). As expected, correlations (i) decrease with decreasing position bias score *b* (comparison within *R* groups) and (ii) decrease with decreasing *R* (comparison within the same position bias level). Hence, the strongest correlations are seen for the most strongly position-biased gene families in species with large *R* values, with median *r*^2^=0.14 for the top 10% and median *r*^2^=0.10 for the top 10%-20%. Interestingly, this analysis also revealed that the most strongly position-biased gene families (top 10%) tend to be small compared to other gene families (**Table S2**).

We repeated this analysis at lower taxonomic ranks, examining correlations across classes, orders, and families within the same higher-level taxonomic rank (**Fig. S5**). Correlations across classes mirror those across phyla, with even stronger correlations especially in the top 10% position-biased genes (**Fig. S5a**). While positional correlations are even stronger for comparisons across orders and families, substantial correlations also appear for gene families without evidence of position biases (**Fig. S5b**,**c**); at this level of evolutionary distance, many positions may be similar simply because of shared ancestry. We thus conclude that gene positioning of strongly position-biased gene families is maintained by selection, and that this selection is strongest in species in which DNA replication times are much longer than cell duplication times (high *R*).

### Expression requirements largely determine position bias levels and preferred positions of genes

With increasing growth rate, a bacterium needs higher expression levels of genes that directly support cell growth, such as those encoding the ribosome, RNA polymerase, and ATP synthase. In contrast, the flagellum does not contribute directly to cell growth, and its required expression decreases with growth rate^17,18^. According to our hypothesis that positional biases evolved under selection for growth rate-dependent gene copy number changes, we expect that genes that directly support growth are biased toward *oriC*, while genes that make no direct contributions to growth should be biased toward *ter*. To test these predictions, we next examined individual functional gene categories, asking which fraction of the corresponding genes tend to have biased positions, and where in the genome these positions are. For this, we divided our normalized genome position scale into three equally sized regions: *oriC* (0–1/3), mid (1/3– 2/3), and *ter* (2/3–1). To reduce noise, we restrict the following analyses to large-*R* species, where gene position biases are strongest, and consider only the top 30% position-biased gene families identified from these species (bias score *b* > 0.4821, *n* = 1195).

As shown in **Fig. 4a**, the most over-represented COG category is cell motility (N), in which 57% gene families are strongly position-biased. Interestingly, this category is only enriched in the *ter* region. At faster growth rates, cells may have a lower incentive to move, but also tend to be larger, so that they likely invest progressively less into motility. The positional biases of flagellar genes, which fall into this category, are detailed in **Fig. 4b**. The second most over-represented COG category is D, which comprises genes related to the cell cycle: of these, 55% are strongly position-biased. This category is enriched not only for gene families biased toward *oriC*, but also for gene families biased toward *ter*. A striking example of cell cycle regulation through gene position is the process of sporulation in *Bacillus subtilis*, which is regulated through the balance between DNA copies of Spo0F, located near *oriC*, and KinA, located near *ter*^8^. Our findings suggest that the positions of many more bacterial genes may contribute to cell cycle control in similar ways.

**Fig. 4.**
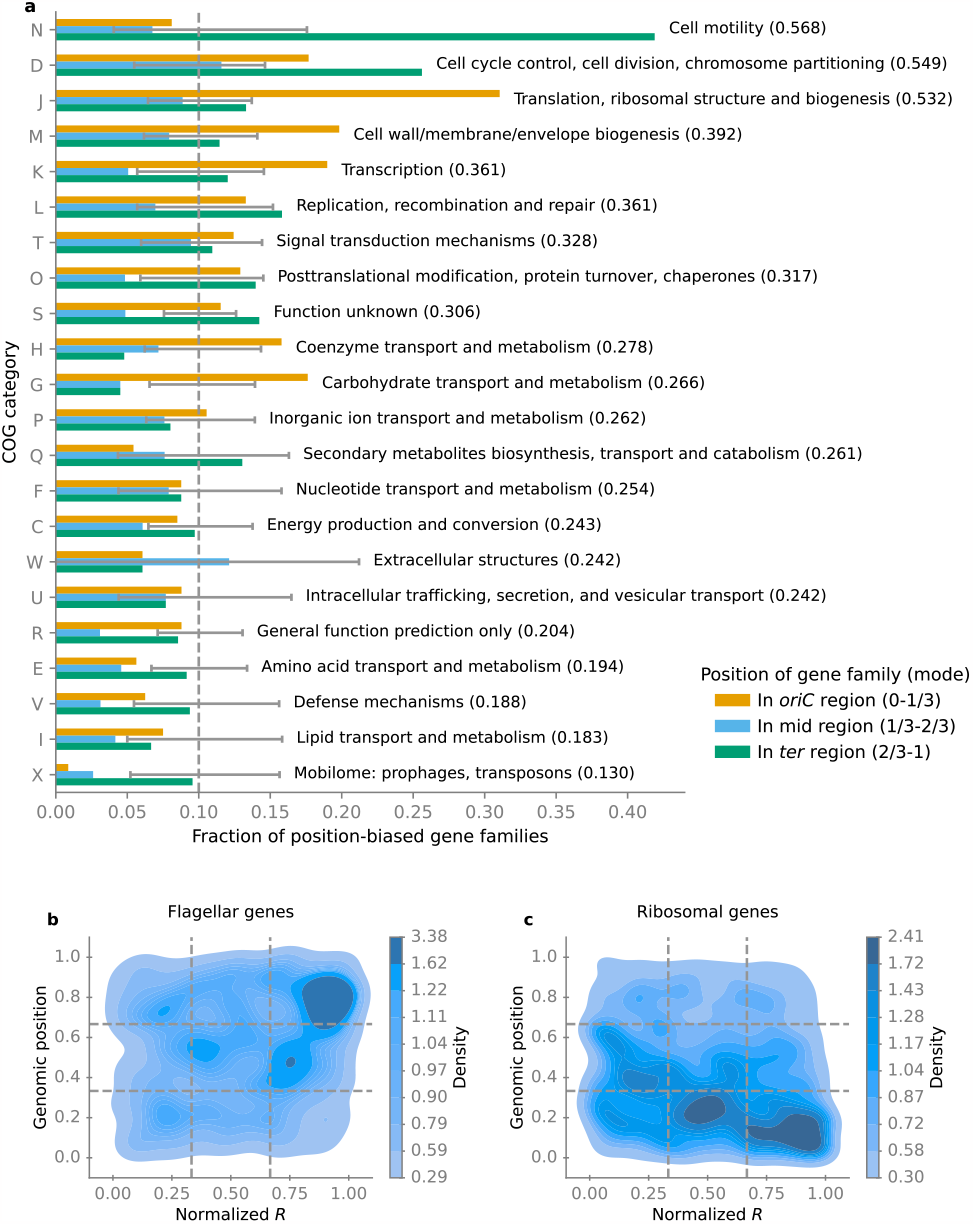
Position biases differ strongly between functional gene categories. (**a**) The 22 COG categories were sorted in descending order of their fraction of strongly position-biased gene families among large*-R* species (top 30%; fractions shown in parentheses). Colored bars split these fractions between the *oriC*, mid, and *ter* regions. If all strongly position-biased genes were distributed randomly across genomes and COG categories, the fractions in each region should all be 0.1 (dashed vertical line); the error bars mark the 95% confidence intervals under this null model. **Fig. S6** shows the same data in terms of absolute numbers rather than fractions. (**b**) Density distribution of flagellar genes and (**c**) density distribution of ribosomal protein genes. To make the plots easier to interpret, we normalized *R* by rank, such that the *R* values are distributed uniformly between the lowest (0) and the largest (1) values observed in our dataset. Each contour level from dark blue to light blue encompasses an additional 10% of the total number of genes in this category. Horizontal lines separate the genomic positions into *oriC*, mid, and *ter* regions. Vertical lines separate species into small-*R*, moderate-*R*, and large-*R* growth groups.

As expected^4^, category J, which comprises translation, ribosomal structure and biogenesis and includes ribosomal genes, is enriched for gene families biased toward *oriC*; however, it is not the most over-represented functional group, despite previous considerations of ribosomal genes as the most – if not the only – positionally biased genes. **Suppl. Text A** discusses the details of positional biases for specific gene families: those encoding ATP synthase, RNA polymerase, murein lipoprotein, DNA topoisomerases, and nucleoid-associated proteins, as well as several other extremely position-biased, large gene families.

Surprisingly, in terms of absolute numbers, the category of gene families with unknown functions contains more strongly position-biased genes than any other functional category, and is enriched for gene families biased to both the *oriC* and the *ter* region (**Fig. S6**). The functions of these genes and the reasons for their positional biases remain to be studied.

In contrast to many other functional categories, the optimal expression of genes encoding metabolic enzymes is usually not a simple function of the cellular growth rate, but depends on specific nutrient conditions. For example, in *E. coli*, when amino acids are supplied as nutrients, the expression of amino acid synthesis pathways drops dramatically compared to minimal carbon media^24^. The environment- and species-specific use of alternative transporters, isoenzymes, and alternative pathways makes any universal growth rate-dependent patterns further unlikely. As expected from these considerations, many of the COG categories that contain fewer position-biased gene families than expected comprise metabolic genes (**Fig. 4a**), including the transport and metabolism of lipids (I), amino acids (E), nucleotides (F), secondary metabolites (Q), inorganic ion (P), coenzymes (H), and carbohydrates (G), as well as energy production and conversion (C).

An exception to this pattern are the cell envelope synthesis pathways in COG category M, which contains a high number of strongly position-biased genes (**Fig. 4a**). Unlike most other metabolic pathways, cell wall synthesis pathways synthesize macromolecules rather than small molecules. While small molecules can either be synthesized by the cell itself or imported from the environment, macromolecules must be synthesized by the cell itself. Moreover, the cellular demand for cell wall synthesis proteins depends on the growth rate; it may hence not be surprising that cell wall synthesis genes have more biased positions than other metabolic genes.

### Genes of related functions can have very different preferred positions

Distinct positional biases of functionally related gene families may reflect specific expression requirements, related to distinct roles in a given cellular process. For example, our recent work found that the different positions of rRNA and tRNA genes help to ensure their optimal relative dosage, facilitating optimal translation efficiency^14^. Another recent work found that the positions of two sporulation genes (Spo0F and KinA) in *B. subtilis* at opposing chromosomal ends regulate the switch between sporulation and vegetative growth^8^. To dig deeper into the role of chromosomal positions in these two processes, we here examine the positions of individual ribosomal genes and sporulation genes in large-*R* species.

Only 6 out of 55 genes encoding ribosomal proteins (gene family size *n*>5) are not positionally biased (**Table S4**). However, although most of the remaining ribosomal genes are biased toward *oriC* (**Fig. 4c, Fig. 5a**), the genes for L20, L32, L35, S1, S16, and L7Ae are systematically located near *ter* (**Fig. 5a**). These contrasting position preferences indicate functional heterogeneity. In particular, ribosomal protein L20 is one of the most important proteins in the early stage of 50S subunit assembly^29^, and its positional bias toward *ter* indicates natural selection for a decrease of its expression with increasing growth rate, warranting further study.

**Fig. 5.**
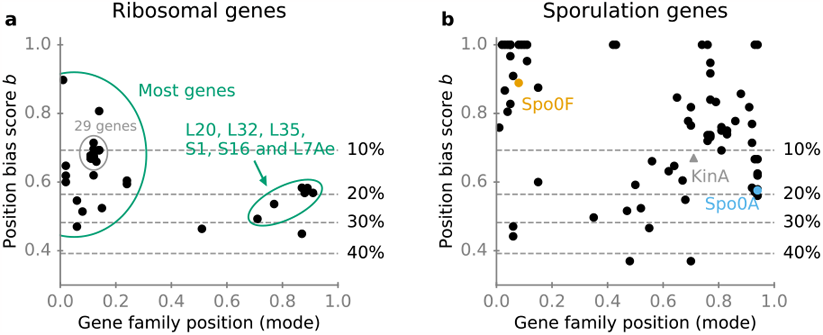
Ribosomal (a) and sporulation (b) genes are biased toward both the origin and the terminus of replication. Horizontal lines from bottom to top mark the threshold of position bias scores at top 40%, 30%, 20%, and 10%, respectively. Genes without statistically significant position biases are not shown, except for KinA, shown as a triangle in panel b.

72 out of 133 sporulation genes (gene family size *n*>5) show positional biases (**Table S4**). Their bias scores are typically even higher than those of ribosomal genes, and they are enriched in both *oriC* and *ter* regions (**Fig. 5b**). For the two sporulation-controlling genes in *B. subtilis*^8^, we found that the position of Spo0F is strongly biased across species, while KinA is slightly preferred at the terminus, although its chromosomal distribution is not significantly different from the uniform distribution (adjusted-*p* = 0.28, *n* = 9) (**Fig. 5b**). Moreover, the sporulation master regulator, Spo0A^30^, is located near *ter* (**Fig. 5b**), consistent with an observed increase of its expression with decreasing growth rate in *B. subtilis*^31^. These findings are consistent with the notion that spores are produced for survival in harsh environments. In addition to the three examples discussed here, there are many more genes involved in sporulation that have biased positions (**Fig. 5b**); these locations may provide hints for their functions, which need to be uncovered by future experiments.

### Growth rate-dependent gene copy number changes do not replace regulation through transcription factors

Most gene regulation in bacteria is believed to be mediated through transcription factors. However, according to our hypothesis, gene families with strong positional biases take care of at least some of their regulatory needs through replication-associated gene dosage effects. One might hence speculate that these gene families require less regulation by transcription factors compared to gene families without positional biases. To test this notion, we compared the fraction of genes known to be regulated through transcription factors in strongly position-biased gene families (top 30%) and all other gene families (**Fig. S7**). We found that strongly position-biased genes are only slightly less frequently regulated by transcription factors than other genes in *E. coli* (52.1% vs 55.6%; *p* = 0.03, one-sided Fisher’s exact test). We did not find the expected systematic differences in *B. subtilis* (37.9% vs 38.7%; *p* = 0.36, one-sided Fisher’s exact test) (**Fig. S7**). These findings indicate that growth rate-dependent regulation through replication-associated gene dosage effects does not replace condition-dependent regulation through transcription factors. Instead, it likely represents an additional layer of regulation. This conclusion is consistent with our previous findings for the chromosomal positions of ribosomal RNA and tRNAs in *E. coli*^14^. The growth rate dependence of their relative copy numbers is not sufficient to result in optimal RNA abundances at lower growth rates without additional transcriptional regulation. However, maximal transcription of all loci results in the optimal relative expression levels at the highest growth rates.

## Conclusions

Previously, chromosomal position biases were known for only a minority of genes, specifically those associated with translation and transcription^4,22^ as well as with sporulation^8,31^. We hypothesized that, more generally, genes might benefit from growth rate-dependent changes in relative copy numbers caused by their specific chromosomal positioning. From this hypothesis, we derived six specific predictions: (1) a significant number of bacterial gene families should show positional biases; (2) position-biased genes should be located predominantly near the origin and near the terminus of replication; (3) with increasing growth rate, protein abundances should increase for genes near the origin and decrease for genes near the terminus of replication; (4) positional biases should be strongest in species whose DNA replication time exceeds the cellular doubling time, and weakest in species for which DNA replication is much faster than cell doubling; (5) the genomic positions of genes from strongly position-biased families should be correlated over vast evolutionary distances; and (6) positional biases should differ in systematic and predictable ways across functional gene categories. All six predictions were fully supported by our analyses. Thus, our results indicate that natural selection for specific growth rate-dependencies of relative gene copy numbers is widespread and has shaped positional preferences for nearly half of all gene families. These findings are consistent with observed evolutionary constraints on chromosomal organization, such as a bias of successful chromosomal inversions toward those that are symmetric around *oriC*^32,33^, thereby conserving the growth rate dependence of replication-associated gene dosage effects.

It has previously been argued that chromosomal biases within *E. coli* are related to the organization of the chromosome into macrodomains and to a gradient of DNA superhelical density from the origin to the terminus of replication^6^. However, the consistency of positional biases across bacterial species suggests a more general underlying mechanistic cause, such as the replication-associated gene dosage effects discussed here.

Chromosomal position biases can provide intriguing clues to gene functions. For example, many genes involved in sporulation are biased either toward *oriC* or toward *ter*, indicating that they may implement similar regulatory mechanisms as those unraveled for Spo0F/KinA^8^. While it was previously thought that ribosomal genes are generally biased toward *oriC*, we find that multiple ribosomal gene families are instead strongly biased toward *ter*. Furthermore, many gene families of unknown function exhibit strong position biases, highlighting their physiological significance across growth rates. Thus, our findings not only reveal an intricate link between bacterial growth, resource allocation, and genome organization, but also provide new clues for the functional characterization of gene families. Moreover, that natural evolution has led to preferred chromosomal positions for almost half of all gene families should motivate the careful consideration of gene positions when designing artificial bacterial genomes.

## Methods

### Species dataset

We obtained the annotation of orthologous bacterial gene families from the 2020 update of the COG (Clusters of Orthologous Genes) database, which covers all bacterial genera that had been sequenced by 2019. With minor exceptions, COG contains only a single genome for each genus^19^. Therefore, this dataset not only provides good coverage of microbial diversity, but also balances the representation of different lineages compared to databases that aim to include all sequenced genomes. To ensure that gene positions can be calculated unambiguously and can be fully compared across genomes, we excluded the following types of genomes: (1) genomes with more than one chromosome; (2) genomes with a linear chromosome; and (3) genomes without an annotation of *oriC* in the DoriC database (version 10)^34^. To reduce biases in the dataset, we restricted our utilization to a single genome per species. As the COG database contains two genomes for the species *E. coli* and *Prochlorococcus marinus*, we specifically employed one genome for each of them (the genome of the *E. coli* K-12 strain and the *P. marinus* MIT 9313 strain). This resulted in a dataset of 773 bacterial genomes (species). While 174 of these genomes have multiple annotated *oriCs*, these are always less than 0.3% of the genome length apart. Different *oriC*s are thus expected to have a negligible effect on gene positions and we randomly selected one of the *oriC*s when multiple *oriC*s were annotated. Lineage information was retrieved from the NCBI taxonomy database^35^ based on the taxonomy ID of the corresponding genome assemblies.

### Gene families

Each Cluster of Orthologous Genes (COG) represents a family of protein-coding genes. To reduce the number of genes erroneously assigned to individual COGs, we increased the stringency of gene assignment, retaining only genes with PSI-BLAST e-values <10^−10^ (instead of the default cutoff 0.001). This step reduced the dataset by 11.6%. As meaningful statistics concerning positional biases are not possible for very small gene families, we excluded COGs with very small numbers of genes (n ≤ 5). This preprocessing resulted in 4430 COGs, which constitute the set of gene families analyzed in our study.

### KEGG orthology database

To verify our results based on the COG database, we used the annotation of orthologous genes from the KEGG orthology (KO) database^36^. For 724 out of the 773 species in our COG dataset, KEGG provided orthologous gene annotations corresponding to the same genome assembly and annotation version. The results using the corresponding KEGG orthology annotations are very similar to those reported in the main text obtained using the COG annotation (**Figures S8**-**S10** and **Table S5**).

### Gene positions

For a given protein-coding gene, we defined its distance to *oriC* as the shortest distance from its midpoint location to *oriC* on the circular chromosome, measured in number of nucleotides. We defined its normalized chromosomal position (position in short) by dividing the distance to *oriC* by half the length of the chromosome. Gene positions thus range from just above 0 (at *oriC*) to just below 1 (at *ter*).

### Relative gene dosage

We used the genome replication model developed by Cooper and Helmstetter^12,13^ to calculate the relative gene dosage at different chromosomal positions in *E. coli* as a function of the growth rate (**Fig. 1c**). We first calculated the average gene dosage (gene copies per cell,^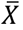^) for a given chromosomal position (*Position*_*x*_),

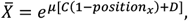

where *μ* is the growth rate, *C* is the DNA replication time (*C* = 41 min in *E. coli*)^12^, and *D* is the time between the termination of one round of DNA replication and the next cell division (*D* = 22 min in *E. coli*)^12^. We then calculated the average genome equivalent of DNA per cell 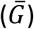,

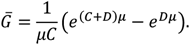

The relative gene dosage is then calculated as the ratio between 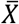 and 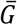.

### Position bias score b

For the normalized chromosomal positions of all genes in a given gene family, we first performed a Gaussian kernel density estimation (bandwidth = 0.05). This procedure provided us with a robust estimate of the mode (distribution peak) of the gene positions. We then drew a sliding window (*width* = 0.3 and *step size* = 0.01) across all chromosomal positions that contained the mode in the window, at each position counting the number of genes covered by the window. We defined the position bias score *b* as the maximal fraction of genes from the gene family covered by one of these windows (see **Fig. S2** for an example).

### Position-biased gene families

We consider a given gene family as *position-biased* if its gene positions across genomes do not follow a uniform distribution. We first performed a Kolmogorov-Smirnov test on each individual gene family against the uniform distribution in the interval [0, 1]. To control the false discovery rate associated with multiple testing, we adjusted the resulting *p*-values using the Benjamini-Hochberg procedure^20^, accepting a false discovery rate of 0.05. Thus, we considered gene families as positional biased if their adjusted *p*-value was below 0.05. Very small gene families could by chance have a high position bias score *b* even in the absence of true positional biases. The Kolmogorov-Smirnov test largely eliminates such cases, without affecting the detection of large position-biased gene families.

We designate the position-biased gene families with the highest position bias scores *b* as *strongly position-biased*, such that the total number of gene families in this subset accounts for 30% of the total gene families in our dataset. The choice of 30% as the cutoff was motivated by the similarity of the distribution of bias scores of this subset to that of ribosomal genes (**Fig. 2b**), which are generally considered to have substantial positional biases.

### Codon adaptation index (CAI) and its normalization

The codon adaptation index (CAI) was calculated according to the original definition^26^, designating ribosomal protein genes as the highly expressed genes. Sequences of ribosomal protein genes were retrieved from the corresponding genomes using the COG annotation.

To facilitate comparisons of CAI across species, we normalized CAI by ranks in each species:

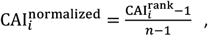

where 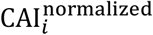 is the normalized CAI of gene *i*, 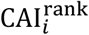 is the CAI rank of gene *i* in the given species in ascending order, and *n* is the total number of protein-coding genes in the given species. The highest CAI in each species was thus normalized to 1, and the lowest CAI was normalized to 0.

### Protein abundance fold-changes

To compare the fold-change in protein abundance of genes encoded in the *oriC*, mid, and *ter* regions, we converted protein abundances (copies per cell in the original datasets) to mass fractions of the total proteome for all three datesets^23–25^. As position-biased gene families tend to be highly expressed (**Fig. 2e**), we only considered genes with relatively high abundances (protein mass fractions greater than 10^−5^ of the total proteome). To be conservative, protein abundance fold-changes greater than 10-fold or less than 0.1-fold (minimal media vs. rich media) were treated as outliers and were excluded from the datasets. We obtained protein abundance fold-changes for 1849 genes from the *E. coli* ribosomal profiling dataset^23^, 1505 genes from the *E. coli* mass spectrometry dataset^24^, and 572 genes from the *B. subtilis* mass spectrometry dataset^25^.

### Maximum growth rate and calculation of R

A previous study collected the maximum growth rate of 214 prokaryotes^22^; 102 of these species are included in our dataset. For all other species in our dataset, we manually searched for scientific publications that report growth rates using Google and Google Scholar with the keywords “doubling time”, “generation time”, and “growth rate” together with the species name (including both the “current” name and “heterotypic synonym” names in the NCBI taxonomy database^35^). In this way, we identified the maximum reported growth rates for an additional 314 species. The final dataset of maximum growth rates for 416 species is listed in **Table S3**, together with the corresponding references (with 102 growth rates taken from Ref.^22^).

We adopted the definition of *R*, the ratio of DNA replication time to the bacterial minimum doubling time, from Ref. 4. Following this publication, we used a constant DNA replication rate of 1000 bp s^-1^. DNA replication times were calculated as half the genome length divided by the DNA replication rate.

To better visualize the density of specific genes along the *R* axis (**Fig. 4** and **Fig. S11-13**), we normalized *R* by rank:

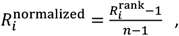

where 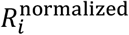 is the normalized *R* of species *i*, 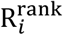 is the *R* rank of species *i* in ascending order, and *n* is the total number of species in the dataset (*n*=416). *R* was thus normalized to a uniform distribution in the interval [0, 1].

### Gene families with multiple genes in one genome

For the purpose of our study, gene duplicates are genes in one genome that are assigned to the same gene family (same COG or KO ID). When calculating position bias scores, the positions of gene duplicates were counted separately. To calculate Pearson’s correlation coefficient of gene positions between two species, we used the arithmetic mean of the positions of all gene duplicates in a given genome to represent the gene family.

### COG membership of ribosomal genes, flagellar genes, and other genes

We labeled gene families as belonging to ribosomal genes, flagellar genes, sporulation genes, RNA polymerase core enzyme, ATP synthase, murein lipoprotein (Lpp), DNA topoisomerases, and nucleoid-associated proteins based on corresponding keyword searches. The COG IDs of these families are listed in **Table S6**.

### Transcriptional regulation datasets

Transcriptional regulation data were downloaded from RegulonDB^37^ for *E. coli*, using only the experimental dataset (version 11.2), and retrieved from Ref.^38^ for *B. subtilis*. Regulation by sigma factors was ignored for our analysis.

### Confidence interval calculation

As a null model for **Fig. 4a**, we assumed that in all COG functional categories, the preferred positions of strongly position-biased genes are uniformly distributed across the *oriC*, mid, and *ter* regions. We defined strongly position-biased genes as those gene families with statistically significant positional biases (at FDR = 0.05) and the highest position bias scores *b*, choosing the *b* cutoff such that they make up 30% of all gene families. Since the *oriC* region contains one-third of all genes, the expected proportion of all gene families in a given functional category that are strongly position-biased toward the *oriC* region is 10% = 30% × ⅓ ; the same is true for the mid and *ter* regions. Let *n* be the total number of gene families in a given COG (KO) category. To obtain a 95%-confidence interval for **Fig. 4a, Fig. S6**, and **Fig. S10**, we drew 10,000 samples from a binomial distribution with *n* trials and 0.1 probability of success. The number of successes was then sorted in ascending order. The 95% confidence interval then ranges from the 250th to the 9750th of the ordered successes.

## Acknowledgments

This work was supported by the Volkswagen Foundation through the “Life?” initiative and by the German Research Foundation (DFG) through grant CRC 1310.

## Author Contributions

XPH designed the study, performed the analyses, and drafted the manuscript. MJL supervised the study. XPH and MJL interpreted the results and edited the manuscript.

