## Supporting Information for "Nearly half of all bacterial gene families are biased toward specific chromosomal positions"

This PDF includes:

- Supplementary Text A
- Supplementary Figures S1-S13
- Supplementary Table S7
- Supplementary References

### Supplementary Text A

#### Positions of genes for ATP synthesis, RNAP polymerase, and Lpp

As expected from our hypothesis that positional biases facilitate growth rate-dependent regulation, the genes encoding RNA polymerase (in category K) and ATP synthase (in category C) tend to be located near *oriC* especially in large-*R* species (**Fig. S11a,b**). With increasing growth rate, replication leads to higher relative gene dosages (more DNA copies) of these genes. The opposite trends are seen for the murein lipoprotein Lpp (COG4238) (**Fig. S11c**), a highly abundant structural protein that links the outer membrane to the peptidoglycan[1,2]. Lpp exhibits the highest CAI among all genes in 17 out of 45 species with this gene. Since bacterial cell sizes increase with growth rate[3,4] and since this increase leads to a decreasing cell surface to volume ratio, the requirement for Lpp also decreases with growth rate; in *E. coli*, for example, the mass fraction of Lpp in the total proteome decreases from 3.7% on minimal medium to 2.5% on rich medium (data from Ref.[5]).

#### Gene positions of DNA topoisomerases and nucleoid-associated proteins (NAPs)

DNA topoisomerases and NAPs do not directly contribute to cell growth. Instead, they affect cell growth by regulating many other genes[6]. In this section, we summarize the positions of genes encoding DNA topoisomerases and some well-studied NAPs in our dataset.

Our dataset contains three frequent DNA topoisomerases: DNA gyrase, DNA topoisomerase IA, and DNA topoisomerase I. We found that gene positions in all of these families are biased both towards *oriC* and towards *ter* (**Fig. S12**). These biases towards opposing chromosomal regions are likely caused by imprecise annotations of these genes. For example, four genes are annotated as DNA gyrase in *E. coli* in COG: DNA gyrase subunit A (GyrA), gyrase subunit B (GyrB), DNA topoisomerase IV subunit A (ParC), and DNA topoisomerase IV subunit B (ParE). While GyrA and GyrB are indeed DNA gyrase, ParC and ParE are subunits of DNA topoisomerase IV, a homolog of DNA gyrase. Despite this potentially noisy annotation, it is evident that DNA topoisomerases, acting as global gene regulators, consistently occupy preferred genomic positions across bacterial species.

Fis is a global transcription regulator of metabolism and ribosomal biosynthesis in the exponential growth phase[7]. Fis tends to be located near *oriC* (**Fig. S13a**). This observation is consistent with the finding that moving Fis from *oriC*-proximal to *ter*-proximal positions reduces cell fitness in *E. coli*[7].

H-NS (which, along with its homolog StpA, is annotated as H-NS in the COG database) is a universal gene expression repressor and increases the survival rate of *E. coli* in stress conditions[8]. Interestingly, while altering the genomic position of H-NS has no short-term impact on fitness in *E. coli*[9], the gene shows a preference for positions near *ter* across species (**Fig. S13b**).

Two other NAPs, IHF and its homolog HU, are annotated as IHF in the COG database. In *E. coli*, IHF[10] and HU[11] play important roles in nucleoid organization and the regulation of genes in response to stress conditions[12]. IHF is located near *ter* (**Fig. S13c**) across species. Again, the minor distribution peak near *oriC* is likely caused by imprecise annotations, as four *E. coli* genes (IhfA, IhfB, HupA, and HupB) are annotated as IHF in the COG database.

Lrp regulates the expression of many metabolic genes[13]. While metabolic genes tend to be less position biased than other genes (**Fig. 4a**), Lrp tends to be located near *oriC* in large-*R* species (**Fig. S13d**).

The MukBEF protein complex (comprising MukB, MukE, and MukF) belongs to the family of SMC (structural maintenance of chromosomes) proteins and is involved in chromosome condensation and partitioning[14]. Remarkably, the genes for all three subunits of MukBEF tend to be located near *ter* across species (**Fig. S13e**). This positioning aligns well with the function of the MukBEF complex, as DNA partitioning occurs after DNA replication finishes, and the *ter*-proximal positioning of MukBEF allows it to double DNA copies shortly before the end of replication, potentially aiding DNA chromosome condensation and partitioning.

Lastly, MatP contributes to the organization of the Ter macrodomain of the *E. coli* chromosome by binding with specific DNA sites (*matS*, a 13 bp motif exclusively found in the Ter macrodomain)[15]. Interestingly, MatP also tends to be located near *ter* (**Fig. S13f**), in close proximity to its target DNA sites.

These examples demonstrate that DNA topoisomerases and NAPs, as global gene regulators in bacteria, share similar genomic positions across species, indicating that their positions may aid in their performance of cellular functions in similar ways.

#### Extremely position-biased large gene families

We also found that some widely distributed gene families (gene family size $n > 500$) exhibit extreme position biases, identifying 13 families with bias scores $b > 0.6$ (**Table S7**). 12 out of the 13 gene families are biased to locations very close to *ori*, while the genes of the last family tend to be located very close to *ter* (COG1559). Two of these gene families encode components of core cellular machineries: a ribosomal protein (COG0230, L34) and a subunit of ATP synthase (COG0712). While the members of the gene families in **Table S7** have highly biased genomic positions in most species, the reasons for these strong biases are not well understood and need to be uncovered by further experimental studies.

#### Table S7. Extremely position-biased large gene families

| Gene family (COG ID) | Gene family size *n* | Mode position | Position bias score *b* | Adjusted *p*-value | COG category | COG Annotation |
|---|---|---|---|---|---|---|
| COG0445 | 656 | 0.01 | 0.833841 | 8.64E-242 | J | tRNA U34 5-carboxymethylaminomethyl modifying enzyme MnmG/GidA |
| COG1195 | 619 | 0.01 | 0.82391 | 1.19E-250 | L | Recombinational DNA repair ATPase RecF |
| COG0592 | 825 | 0.01 | 0.815758 | 0 | L | DNA polymerase III sliding clamp (beta) subunit, PCNA homolog |
| COG0594 | 547 | 0.01 | 0.815356 | 9.20E-212 | J | RNase P protein component |
| COG0357 | 745 | 0.01 | 0.806711 | 9.09E-253 | J | 16S rRNA G527 N7-methylase RsmG (former glucose-inhibited division protein B) |
| COG0230 | 557 | 0.01 | 0.780969 | 1.54E-179 | J | Ribosomal protein L34 |
| COG0486 | 670 | 0.01 | 0.744776 | 6.63E-190 | J | tRNA U34 5-carboxymethylaminomethyl modifying GTPase MnmE/TrmE |
| COG0706 | 869 | 0.01 | 0.69275 | 8.95E-183 | M | Membrane protein insertase Oxa1/YidC/SpoIIIJ |
| COG1475 | 1033 | 0.01 | 0.656341 | 1.22E-189 | D | Chromosome segregation protein Spo0J, contains ParB-like nuclease domain |
| COG0759 | 751 | 0.01 | 0.61518 | 1.23E-114 | M | Membrane-anchored protein YidD, putatitve component of membrane protein insertase Oxa1/YidC/SpoIIIJ |
| COG1559 | 686 | 0.92 | 0.612245 | 1.19E-63 | M | Endolytic transglycosylase MltG, terminates peptidoglycan polymerization |
| COG0593 | 1088 | 0 | 0.610294 | 9.70E-228 | L | Chromosomal replication initiation ATPase DnaA |
| COG0712 | 712 | 0.04 | 0.608146 | 2.48E-59 | C | FoF1-type ATP synthase, delta subunit |

#### Supplementary Figures

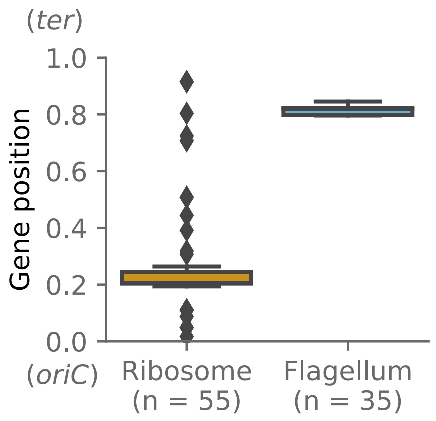


**Fig. S1. Gene positions of ribosomal protein genes and flagellar genes in *E. coli*.** Lines within boxes mark medians. Boxes extend from the first quartile to the third quantile of the data, whiskers extend from the boxes by 1.5x the interquartile range, diamonds are outliers.

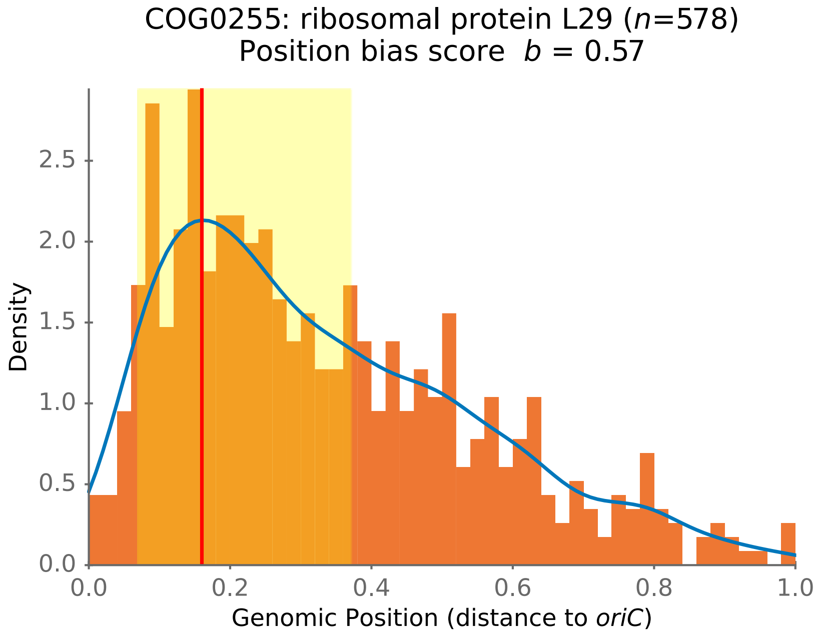


**Fig. S2. Illustration of the definition of position bias score *b*, using COG0255 as an example.** The histogram shows the density of gene positions of COG0255 along the chromosome. The blue curve marks the Gaussian kernel density estimate. The distribution peak (mode) of the gene positions, shown as a vertical red line, is estimated as the maximum of the blue curve. A sliding window of width 0.3 and step size 0.01 is used to find the maximal fraction of genes located in such a window around the estimated mode. The sliding window with most genes is shown in light yellow. The fraction of genes located in this window is defined as the position bias score *b* (*b* = 0.57 in this example).

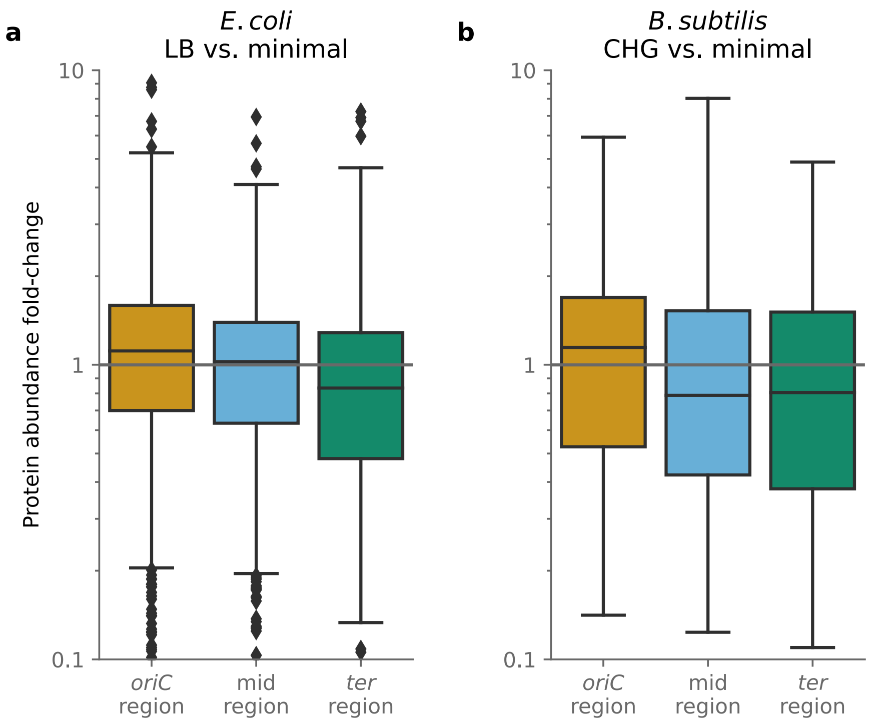


**Fig. S3. Fold-change of protein abundance of genes located in the *oriC*, mid, and *ter* regions.** (**a**) Protein abundance on LB media vs. the averaged protein abundance on minimal media, quantified by mass spectrometry in *E. coli*. Data from Ref.[16]. (**b**) Protein abundance on CHG media (amino acid-based medium + glucose) vs. the averaged protein abundance on minimal media, quantified by mass spectrometry in *B. subtilis*. Data from Ref.[17]. Genes biased to the *oriC* region tend to have higher abundances on rich media than on minimal media in both *E. coli* and *B. subtilis* (one-sided Wilcoxon signed-rank tests: $p = 5.2\times10^{-5}$ for *E. coli* ($n = 551$); $p = 3.3\times10^{-3}$ for *B. subtilis* ($n = 186$)). In contrast, genes biased to the *ter* region tend to have higher abundances on minimal media than on rich media (one-sided Wilcoxon signed-rank tests: $p = 5.5\times10^{-8}$ for *E. coli* ($n = 452$); $p = 0.03$ for *B. subtilis* ($n = 196$)). Lines within boxes mark medians. Boxes extend from the first quartile to the third quantile of the data, whiskers extend from the boxes by 1.5x the interquartile range, diamonds are outliers.

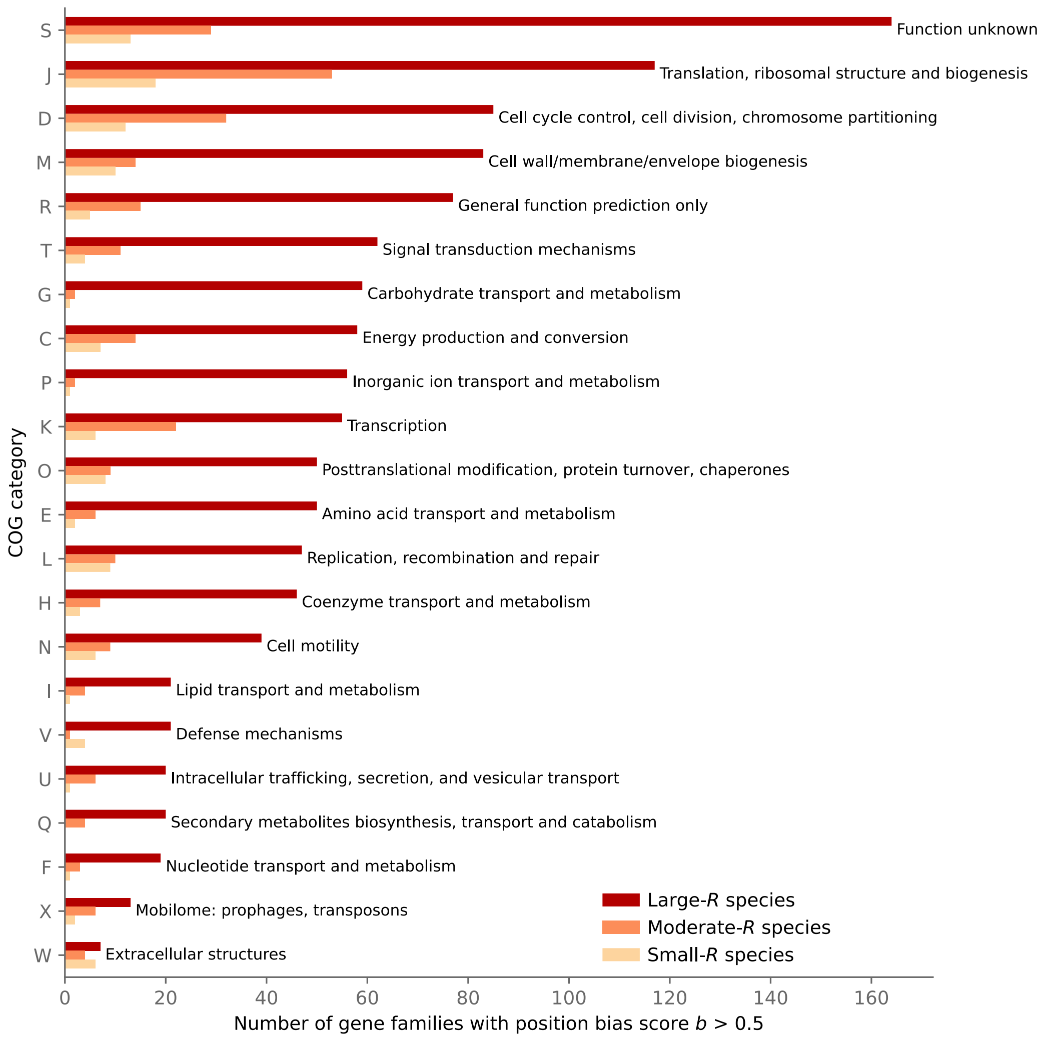


**Fig. S4. Number of gene families with position bias scores greater than 0.5 in different COG catagories in large-*R*, moderate-*R*, and small-*R* growth groups.** The COG categories are sorted in descending order of the number of position-biased gene families in large-R species.

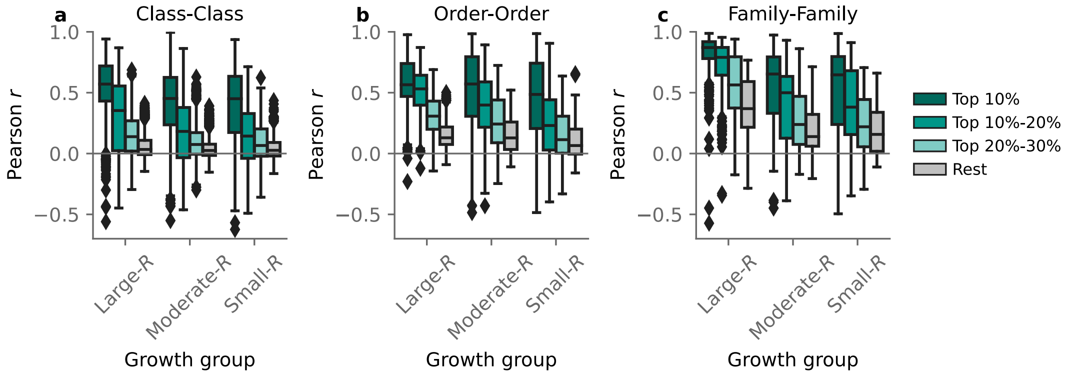


**Fig. S5. Maximal growth rate strongly influences the correlation of gene positions across species.** Distributions of Pearson correlation coefficients between gene positions of (**a**) two species from the same phylum but from two different classes; (**b**) two species from the same class but from two different orders; (**c**) two species from the same order but from two different families. Distributions are shown separately for different position bias levels, which are marked by colors. Lines within boxes mark medians. Boxes extend from the first quartile to the third quantile of the data, whiskers extend from the boxes by 1.5x the interquartile range, diamonds are outliers.

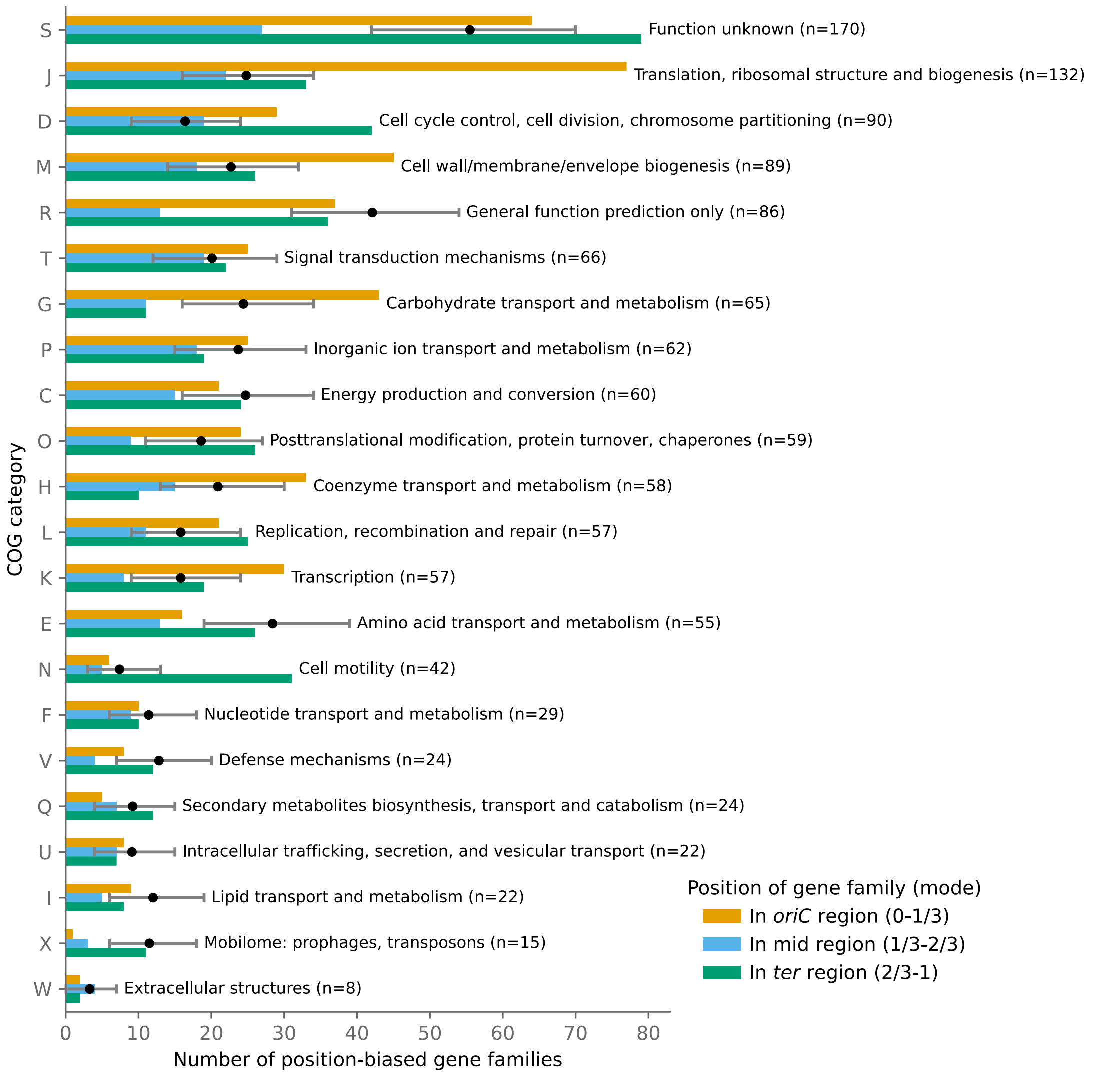


**Fig. S6. The number of gene families that are strongly position-biased differs across COG categories.** The 22 COG categories were sorted in descending order of their total numbers of strongly position-biased gene families among large-*R* species (top 30%; gene family numbers shown in parentheses). Colored bars split these fractions between the *oriC*, mid, and *ter* regions. If all strongly position-biased genes were distributed randomly across genomes and COG categories, the number of strongly position-biased gene families in each region should all correspond to 10% of the total genes in the corresponding COG category (marked by dots; the error bars mark the 95% confidence intervals for expected gene family numbers under this null model). **Fig. 4** in the main text shows the same data in terms of fractions of gene families rather than absolute numbers.

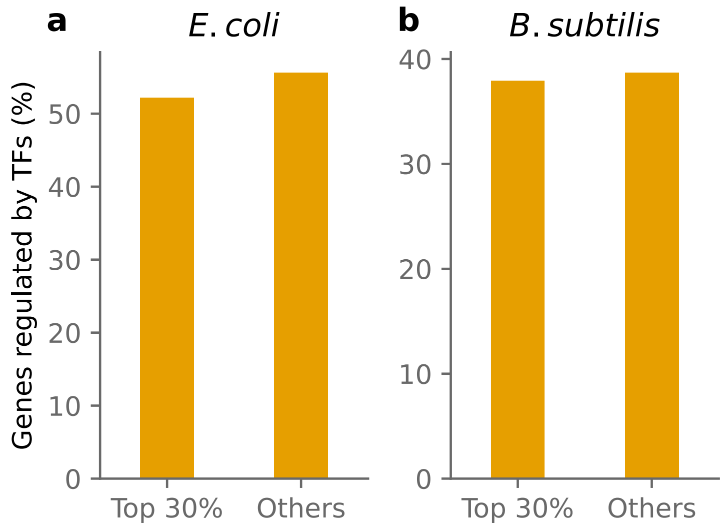


**Fig. S7. Genes regulated by transcription factors (without sigma factors) in (a) *E. coli* and (b) *B. subtilis* at different position bias levels.**

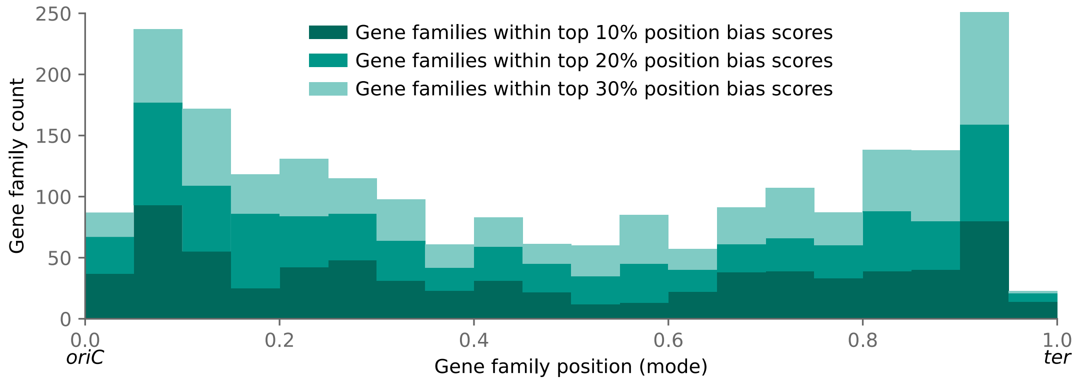


**Fig. S8. Distribution of the mode (gene position peak) of strongly position-biased gene families in the KEGG Orthology (KO) database.** Colors represent position bias levels.

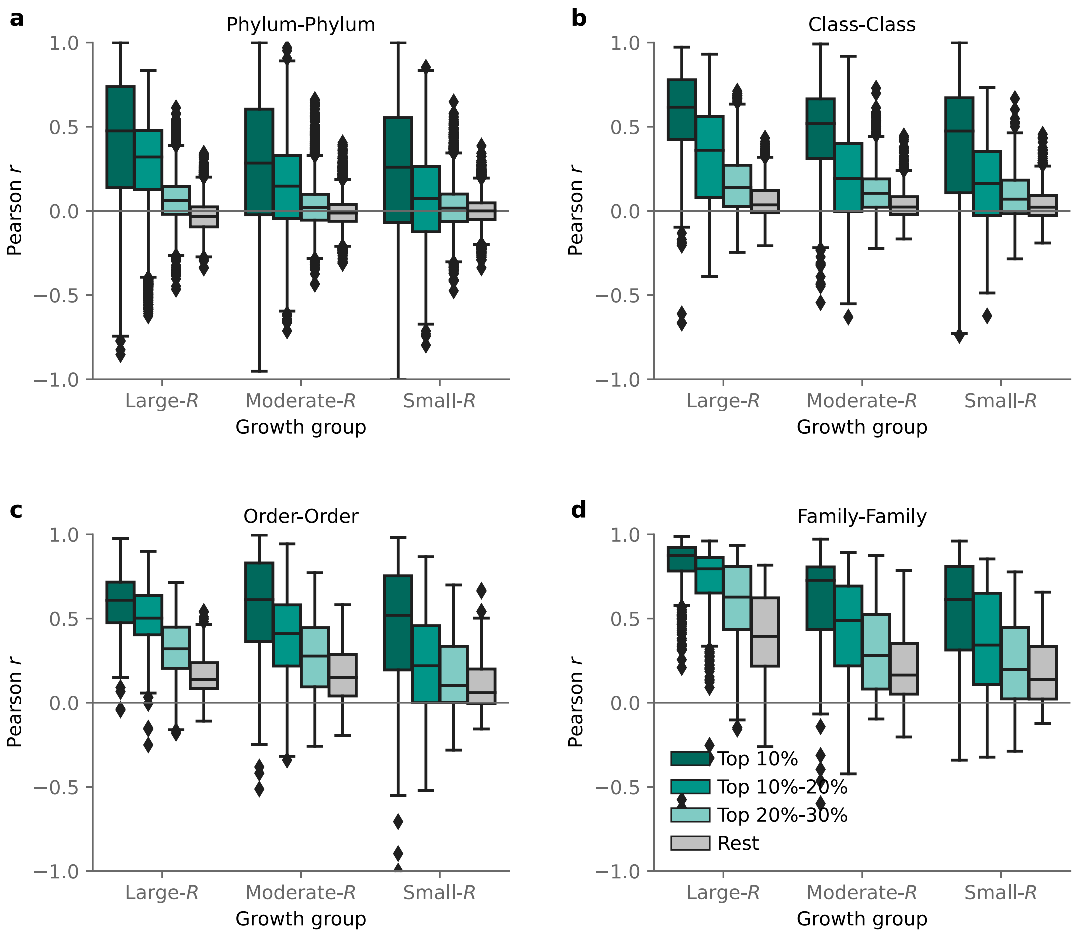


**Fig. S9. Maximal growth rate strongly influnces the correlations of gene positions across species based on orthologous genes annotations from the KEGG orthology (KO) database.** Distributions of Pearson correlation coefficients between gene positions of (**a**) two species from different phyla, (**b**) two species from the same phylum but from different classes, (**c**) two species from the same class but from different orders, and (**d**) two species from the same order but from different families. Distributions are shown sepearately for different position bias levels, which are marked by colors. Lines within boxes mark medians. Boxes extend from the first quartile to the third quantile of the data, whiskers extend from the boxes by 1.5x the interquartile range, diamonds are outliers.

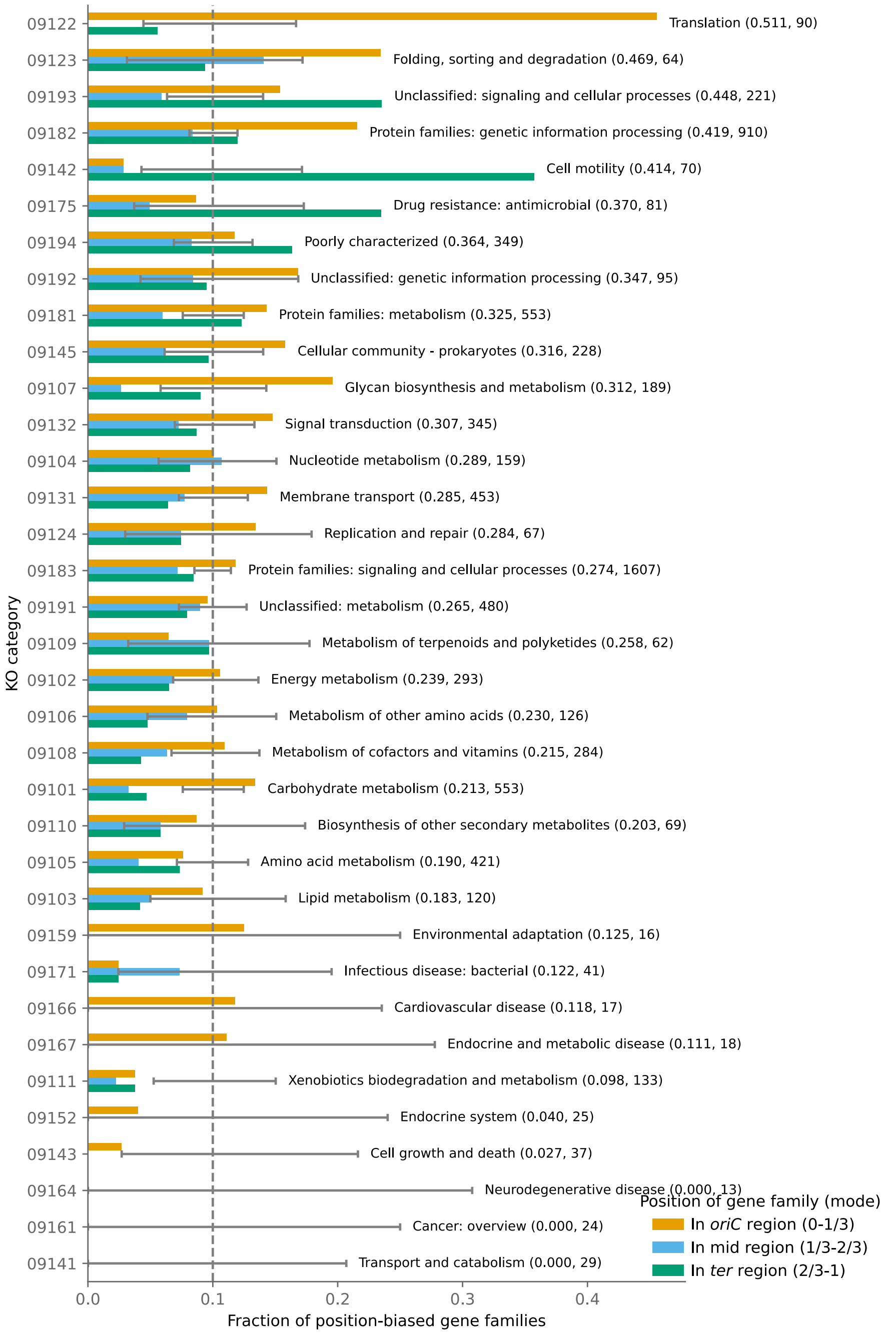


**Fig. S10. Position biases differ strongly between functional gene categories as annotated in the KEGG orthology (KO) database.** The 35 KO categories were sorted in descending order of their fraction of strongly position-biased gene families among large-*R* species (top 30%; fractions shown as the first number in parentheses). The second number in parentheses shows the total number of gene families in the corresponding KO category. Colored bars split these fractions between the *oriC*, mid, and *ter* regions. If all strongly position-biased genes were distributed randomly across genomes and KO categories, the fractions in each region should all be 0.1 (dashed vertical line); the error bars mark the 95% confidence intervals under this null model. The KO categories on the y-axis are IDs in the secondary hierarchy of KEGG Brite tables for KEGG Orthology (extracted from https://www.kegg.jp/brite/ko00001).

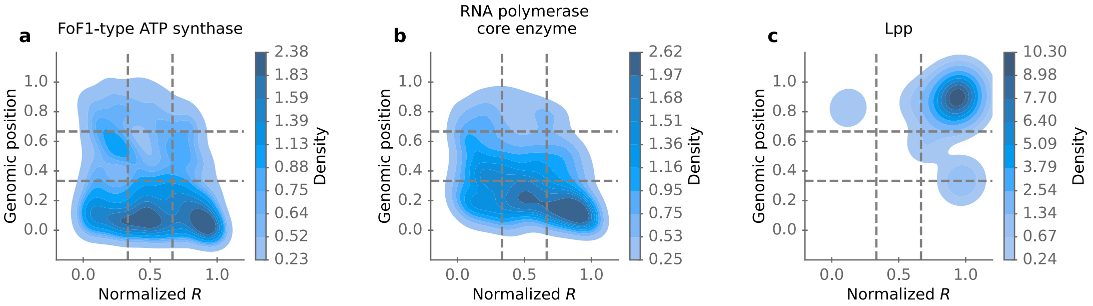


**Fig. S11. Density distributions of (a) FoF1-type ATP synthase genes, (b) RNA polymerase core enzyme, and (c) murein lipoprotein.** To make the plots easier to interpret, we normalized *R* by rank, such that the *R* values are distributed uniformly between the lowest (0) and the largest (1) values observed in our dataset. Each contour level from dark blue to light blue contains an additional 10% of the total number of genes in this category. Horizontal lines separate the genomic positions into *oriC*, mid, and *ter* regions. Vertical lines separate species into small-*R*, moderate-*R*, and large-*R* growth groups.

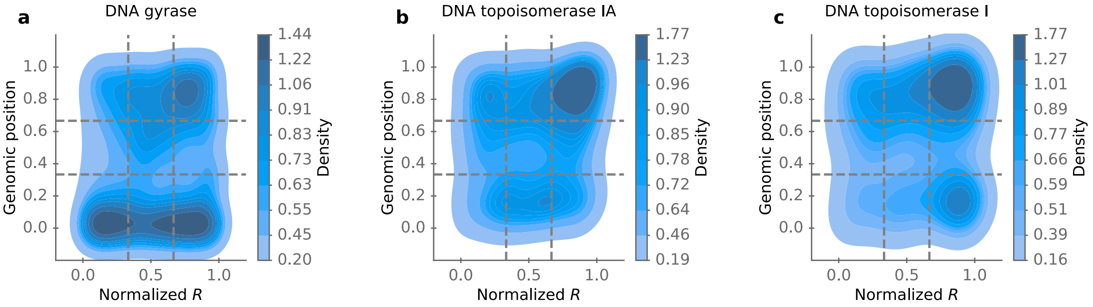


**Fig. S12. Density distributions of DNA topoisomerases.** (**a**) DNA gyrase, (**b**) DNA topoisomerase IA, and (**c**) DNA topoisomerase I. To make the plots easier to interpret, we normalized *R* by rank, such that the *R* values are distributed uniformly between the lowest (0) and the largest (1) values observed in our dataset. Each contour level from dark blue to light blue contains an additional 10% of the total number of genes in this category. Horizontal lines separate the genomic positions into *oriC*, mid, and *ter* regions. Vertical lines separate species into small-*R*, moderate-*R*, and large-*R* growth groups.

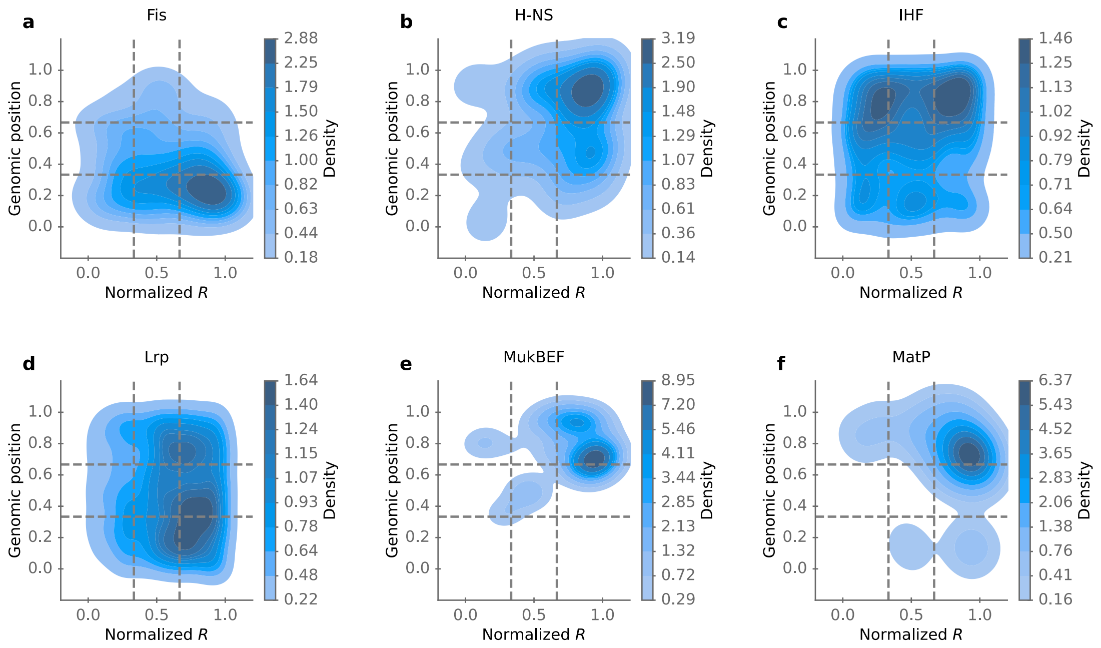


**Fig. S13. Density distributions of nucleoid-associated proteins.** (**a**) Fis, (**b**) H-NS, (**c**) IHF, (**d**) Lrp, (**e**)MukBEF, and (**f**) MatP. To make the plots easier to interpret, we normalized *R* by rank, such that the *R* values are distributed uniformly between the lowest (0) and the largest (1) values observed in our dataset. Each contour level from dark blue to light blue contains an additional 10% of the total number of genes in this category. Horizontal lines separate the genomic positions into *oriC*, mid, and *ter* regions. Vertical lines separate species into small-*R*, moderate-*R*, and large-*R* growth groups.
